# TGF-β1-induced differentiation enhances chemotherapy response in metastatic colorectal cancer organoids

**DOI:** 10.64898/2026.06.04.730067

**Authors:** Sabrina J. Fletcher, Letizia Pizzini, Irene Catalano, Miriam Palmiero, Federica Galvagno, Sofia Borgato, Elena Grassi, Andrea Bertotti, Luca Primo, Livio Trusolino, Alberto Puliafito

## Abstract

**Background:** In metastatic colorectal cancer, systemic therapies frequently fail, partly due to underlying phenotypic plasticity rooted in pre-existing multi-type cell populations. Intratumoral lineage hierarchies within colorectal tumors require renovated efforts to decode growth principles, design rational therapeutic approaches, and accurately interpret drug response. Understanding the cell-state dynamics of untreated tumors and the degree of cell responsiveness to exogenous stimuli is therefore crucial to improving currently underwhelming therapeutic outcomes.

**Methods:** Here, we leveraged patient-derived organoids established from hepatic metastases of colorectal cancer patients to deconstruct population hierarchies by combining single-cell transcriptomics with single-molecule RNA fluorescent in situ hybridization. Computational frameworks were used to identify independent gene modules. We then employed flow cytometry analysis to track cytokine-induced population shifts using a cell surface marker, validating our findings through bulk RNA analysis, functional assays, and viability assays in response to oxaliplatin.

**Results:** Our data substantiate the existence of a dual population configuration within untreated metastatic colorectal cancer organoids with diverse genetic backgrounds. Gene modules detected via single-cell transcriptomics delineate a stem-like (LGR5+) and a differentiated-like (KRT20+) population that fluctuate dynamically over time. Spatially and temporally resolved, single-cell level analysis through single molecule FISH captures the inherent stochasticity in cell fate decisions revealing surprising phenotypic variability even across different organoids derived from the same patient. By using GABRA2 as a surface marker we track the emergence of differentiated cells over the course of time and investigate the respective roles of TGF-β1 and IL-6 in the differentiation of organoids. Our findings indicate that IL-6 exerts no major effect within our cell autonomous setting. In stark contrast, TGF-β1 triggered cell cycle arrest and differentiation, while simultaneously reducing clonogenic capacity and significantly amplifying the cytotoxic potency of oxaliplatin.

**Conclusions:** Our findings provide evidence of the dual Stem and Differentiated population hierarchy in metastatic colorectal cancer organoids, and demonstrate how this axis can be effectively hijacked by TGF-β1 to suppress tumor growth. Our results suggest that further mechanistic exploitation of this cell-autonomous, tumor-suppressive arm of TGF-β1 signalling could open unappreciated therapeutic windows in advanced colorectal cancer.

## INTRODUCTION

The intestine is a highly organized tissue characterized by a well-defined cellular hierarchy (1, 2). Intestinal homeostasis and regeneration are ensured by tightly controlled transitions between different composing cell types and a selective phenotypic plasticity. Interestingly, a similar hierarchical organization is observed in colorectal tumors. Evidence supports the presence of cancer stem-like cells, considered to be associated with the development of colorectal cancer (CRC), alongside cells exhibiting varying degrees of differentiation, recapitulating features of the normal intestinal epithelium (2–4).

Notably, several molecular and phenotypic features are shared between cell types found in the normal intestine and in tumors; for example, both intestinal stem cells (ISCs) and cancer stem-like cells have been associated with high expression of the known markers LGR5 and EPHB2 and are equipped with self-renewal capacity and clonogenic potential (3, 5). Likewise, differentiated cells, although phenotypically diverse, exhibit shared characteristics, including KRT20 expression along with reduced cell cycle activity (6–9).

Such pronounced intratumor phenotypic heterogeneity is manifested in the emergence of distinct subpopulations within tumors, which have been associated with tumor development, progression, and variable response to treatment (10–15). Different subpopulations, indeed, respond differentially to therapies and therefore efforts have been made to understand which cell-state transitions could be mechanistically controlled to enhance treatment response. For example, chemotherapy and radiotherapy primarily target highly proliferative cells. In theory, a higher proportion of proliferative, stem-like cells within a tumor could improve relative treatment efficacy. However, the ability of these cells to exist in or enter a quiescent state has been linked to treatment resistance (15–17). Conversely, inducing a shift towards differentiation - an approach known as differentiation therapy – is aimed at reducing the proportion of proliferative cells while increasing the fraction of differentiated cells, which are generally considered to be non-proliferative and prone to cell death (18–20). Such a clear conceptual approach is however hindered by the reported plasticity of many of the subpopulations characterized in colorectal cancer. For example, LGR5 expressing cells have been shown to be associated to a slow-cycling subpopulation, associated to chemoresistance (15, 21). The same holds for KRT20-expressing cells, associated with a differentiated state, which however retain the ability to revert to a stem-like state and re-express LGR5 under specific conditions, being therefore more dynamic and less terminally differentiated than previously thought (4).

The observed phenotypic plasticity in colorectal cancer partly explains the reasons of systemic therapy failure and the emergence of minimal residual disease in the context of metastatic disease (22). Given the limited therapeutic options in this context, discovering actionable molecular switches that might imbalance the population towards specific states could represent a valuable option to steer the population into a more vulnerable condition for the current therapy.

In order to dissect the dynamic transitions between cellular states, it is necessary to resort to model systems that can faithfully recapitulate both cellular heterogeneity and plasticity. In this respect, patient-derived organoids have several key advantages over other biological models and have been successfully used to model the normal intestine, related diseases and cancer (23–27). Specifically, tumor organoids have the ability to develop from single cells into complex three-dimensional structures enabling the recapitulation of key features of cancerous tissue organization and, in particular, as cells proliferate to form organoid structures, they can acquire distinctive differentiation states. In addition, organoids recapitulate spatial organization and apico-basal polarization, further reflecting features of the tissue of origin (28). The resulting cellular heterogeneity and hierarchical organization make organoids an ideal model for studying intratumor phenotypic heterogeneity (29, 30).

Leveraging the organoid culture system, we set out to examine the cellular hierarchy and intratumor phenotypic heterogeneity within colorectal tumors. To this aim, we used a set of organoids from the XENTURION biobank, which were originally derived from patient-derived xenografts (PDXs) of CRC liver metastases (31). We complemented this with single-cell techniques, including single-cell RNA sequencing (scRNAseq) and fluorescent in situ hybridization (FISH), to identify transcriptionally distinct subpopulations in genetically distinct organoids. Furthermore, we sought to modulate the proportions of each subpopulation through exogenous stimulation and ultimately evaluate the consequent impact on treatment response.

## METHODS

### Organoid culture

Organoids were obtained from the XENTURION biobank of PDX-derived mCRC organoids developed at Candiolo Cancer Institute (31). For regular culturing, they were embedded in Matrigel® Growth Factor Reduced (GFR) Basement Membrane Matrix (Corning) or Cultrex Basement Membrane Extract (BME Type 2 or Ultimatrix RGF BME, R&D Systems). The complete growth medium consisted of Dulbecco’s modified Eagle medium/F12 (DMEM/F12) supplemented with penicillin-streptomycin, 2 mM L-glutamine, 1 x B27 (Gibco), 1 x N-2 supplement (Gibco), 1 mM N-acetylcysteine (NAC), and 20 ng/ml of human EGF (Sigma-Aldrich). Organoids were cultured at 37°C in a humidified atmosphere with 5% CO2, and were tested to exclude mycoplasma contamination.

### Single-cell RNA sequencing (scRNAseq) – Samples preparation, library generation and sequencing

Organoid cells were plated as single cell in 12-well plates and cultured in complete growth medium for the indicated number of days. Organoids were then collected by centrifugation, dissociated into single cells using 0.05% Trypsin-EDTA (Gibco), washed, stained with DAPI, washed again, and resuspended in PBS containing 0.05% Bovine Serum Albumin (BSA). The cell suspension was passed through a 40 µm cell strainer and cells were counted using Trypan Blue (Invitrogen) to exclude dead cells. To obtain a pure, viable single-cell suspension, cells were then sorted using a MoFlo Astrios EQ cell sorter, gating for single cells and excluding dead cells based on DAPI staining. Sorted cells were collected in 0.05% BSA, centrifuged, and counted. The cell suspension was then adjusted to a concentration of 750-1000 cells/μl. In order to analyze 3’ gene expression at single-cell resolution, single-cell barcoding and library construction were performed using the Chromium Next GEM Single Cell 3’ Reagent Kits v3.1 Dual Index and the Chromium Controller instrument (10x Genomics, Inc., Pleasanton, CA, USA), following the manufacturer’s instructions. Briefly, after sorting, the appropriate volume of cell suspension (based on the 10x Genomics user guide) was used to achieve a target recovery of 4000 cells. For both cDNA amplification and indexing PCR steps, 12 cycles were identified as the optimal number under our conditions. Final libraries were quantified using the Qubit dsDNA HS Assay Kit (Thermo Fisher Scientific, Waltham, MA, USA), and their fragment distribution was evaluated using the High-Sensitivity DNA Assay Kit (Agilent Technologies, Santa Clara, CA, USA). Equal amounts of DNA libraries were pooled and sequenced on the NovaSeq 6000 sequencer (Illumina Inc., San Diego, CA, USA), generating sequencing data according to the manufacturer’s settings (read 1: 28 bp; read 2: 90 bp; dual index: 10 bp).

### Single-cell RNA sequencing (scRNAseq) – Computational analysis

Gene counts matrices were obtained for 10x derived samples with Cell Ranger (v3.1.0, reference transcriptome vGRCh38-1.2.0). Downstream analyses were performed using the single-cell analysis suite rCASC (v5.5.9) (32). Quality control procedures were applied to identify and exclude low-quality cells based on ribosomal and mitochondrial gene content, in order to remove cells likely affected by stress or damage during single-cell sorting prior to Chromium processing (scannobyGtf parameters: riboStart.percentage = 1, riboEnd.percentage = 50, mitoStart.percentage = 1, mitoEnd.percentage = 40, thresholdGenes = 250). Ribosomal and mitochondrial genes were removed from the analysis, together with genes showing zero expression in all cells. To improve the reliability of gene expression estimates, data were denoised using SAVER (v1.1.2) (33). The resulting expression matrix, comprising approximately 3,000 cells and 15,000 genes per sample, contained log2-transformed counts per million (CPM) and was subsequently used for downstream analyses and visualization on plots. The gene modules were identified independently for each dataset using Hotspot package with negative binomial model and a k-nearest neighbor graph (k=30) computed in UMAP latent space (34). Genes with significant autocorrelation (FDR < 0.05) were ranked according to their Z-score, and the top 500 genes were used to compute pairwise local correlations. Gene modules were subsequently identified from a gene-gene affinity matrix, using the Hotspot module detection framework, with min genes threshold = 40, core only = False and fdr threshold = 0.05. This procedure identified all the gene modules active in each dataset. For each dataset, two gene modules associated with known stem-like and differentiated-like transcriptional programs were identified. In CRC0327 at 11 dps, the Stem module was obtained by the union of two modules, one more enriched in cells in S and G2/M phases and the other with cells in G1, both contributing to the Stem transcriptional program. In addition, one or more gene modules displaying proliferative characteristics were identified in each dataset. These clearly overlapped with the cell populations assigned to G2/M and S phases by cell cycle scoring analysis (see below). The final modules used in this study are summarized in Supplementary Table 1. To identify stem- and differentiated-like cell populations, Leiden clustering was performed independently on each dataset using only the genes belonging to the corresponding stem or differentiated module. Dimensionality reduction and neighborhood graph construction, prior to clustering, were computed using Scanpy (35). Clustering granularity was selected based on the average silhouette score computed across all cells, evaluated over multiple resolutions, and choosing the non-zero resolution yielding the highest silhouette values. For each module, we computed a single-cell metagene value by calculating the geometric mean of the genes within the module. Cell populations associated to Stem and Differentiated modules were then selected by picking those with the maximum module metagene score. From the integration of stem and differentiated module–based classifications, we obtained cells that were positive or negative to both calls. These were identified as double-positive and double-negative cells.

Throughout the paper, we refer to “minimal modules” (Supplementary Table 2) as the two gene sets defining each phenotype (Stem and Differentiated), obtained as the intersection of the corresponding modules identified across the three patients at 11 dps. Parallel to the module-based analyses, cell cycle states were inferred for each dataset using established S-phase and G2/M-phase scoring procedures. Cell cycle scoring was performed using Scanpy (35).

### XENTURION biobank expression analysis

Publicly available bulk RNA-seq data of CRC liver metastasis organoids from the XENTURION biobank (EGA accession: EGAS00001007024) were used to compare expression from our samples to the full biobank cohort based on minimal modules expression profiles (31). Five of the 220 samples, corresponding to three original models, were originally excluded because of concerns regarding specimen identity raised by transcriptional analyses or post hoc pathological revision (31). Replicate samples corresponding to the same patient were averaged to obtain a single patient-level expression profile (136 samples), followed by log_2_ transformation. Metagene scores were computed for both Stem and Differentiated minimal modules as the geometric mean of the expression levels of genes belonging to each signature. Scores were subsequently standardized using z-score normalization. For Figure 3G, the differentiated module was modified by excluding GABRA2 prior to metagene computation, to enable direct comparison with GABRA2 z-score across all samples.

### Single-molecule Fluorescence In Situ Hybridization (smFISH)

For time course experiments, organoids were plated as single cells in 12-well plates, cultured for the indicated number of days, and then collected by centrifugation. For cytokine treatments, organoids were cultured for 11 days, after which human IL-6 (50 ng/mL; #130-093-931, Miltenyi Biotec) or human TGF-beta 1 Recombinant Protein (5 ng/mL; #100-21-10UG, PeproTech®) was added for 3 days prior to collection. To produce Formalin-Fixed, Paraffin-Embedded (FFPE) samples, pellets were resuspended in 1% agarose (#50100, SeaPlaque® Agarose, Lonza). Samples were then fixed for 24 h in 4% paraformaldehyde (PFA; sc-281692, ChemCruz) at 4°C and subsequently embedded in paraffin. Sections of 5 µm thickness were obtained using a microtome. RNA expression of target genes was evaluated on 5 µm FFPE sections, using RNAScope Multiplex Fluorescent Reagent Kit v2 (#323100, Advanced Cell Diagnostics, Bio-techne®), following the manufacturer’s instructions. Briefly, after deparaffinization and peroxidase blocking, samples were pre-treated with boiling antigen retrieval for 15 min and with protease Plus for 30 min at 40°C. Then, samples were incubated for 2 h at 40°C with appropriate mixed probes. The signals were amplified and finally detected with OPAL dyes (1:1500, Aurogene) according to the ACDBio protocol and assigning OPAL520 (#FP1487001KT) at channel 1, OPAL690 (#FP1497001KT) at channel 2, and OPAL570 (#FP1488001KT) at channel 3. Target genes were LGR5 (Hs-LGR5-C3, #311021-C3), Ki67 (Hs-MKI67, #591771), and KRT20 (Hs-KRT20-C2, #549681-C2). The sequences of target probes, preamplifier, amplifier, and label probes are proprietary to the manufacturer (Advanced Cell Diagnostics, Hayward, CA). Images of 0.3μm spaced z-stacks were acquired with an inverted wide-field microscope Nikon Ti2 (Lipsi) over the thickness of the slice with a 63X 1.2 NA immersion objective and a digital camera (IRIS 15) and collapsed onto 2D images with either maximum projection or Extended Depth of Focus (for fluorescence or bright-field respectively). *Image analysis*: In order to quantify the expression of each transcript, we reconstructed cell regions of interest (ROI) to assign each spot to a given cell. We first segmented nuclei using Cellpose (36) and computed a nuclear labelled mask. Cellular ROI were reconstructed by morphological expansion of nuclear masks within organoid boundaries obtained from segmentation of bright-field images, in order to prevent label propagation outside the organoid area. Spot detection and quantification were performed using a custom Python-based image analysis pipeline. Briefly, maximum-intensity projected smFISH images from each fluorescence channel were sharpened using an unsharp masking filter (radius = 0.56 μm / 4 pixels, amount = 2) to enhance spot signal. Images were then segmented using a three-class multi-level Otsu thresholding approach, retaining only the highest-intensity class to identify candidate RNA spots. To estimate the characteristic size of single RNA spots, we used the mode of the size distribution of segmented objects as a reference single-spot size, and normalized the size of each segmented component accordingly to estimate the corresponding number of RNA spots. The number of spots belonging to each cell in an organoid was then fed to a hierarchical clustering algorithm with four classes, using the number of spots corresponding to LGR5 and KRT20 as features. The same clustering approach was also used to classify cells as MKI67-positive or -negative, using the number of MKI67 spots per cell as the sole feature.

For visualization purposes, single-channel images shown here were autocontrasted with FiJi software (which trims 0.175% of the histogram on the low and high side) (37). Composite images were manually adjusted in order to preserve intensity in all the channels.

### Gene expression analysis of organoids (RT-qPCR)

Organoids were cultured from single cells for the indicated number of days and collected by centrifugation. RNA was extracted using the Maxwell® RSC miRNA Tissue Kit (#AS1460, Promega), according to the manufacturer’s protocol, and RNA concentration was measured using a DS-11+ spectrophotometer (DeNovix Inc.). RNA was treated with RQ1 DNase (#M6101, Promega) and subsequently reverse-transcribed using the High-Capacity cDNA Reverse Transcription Kit (#4368814, Thermo Fisher Scientific) to produce cDNA. No-reverse transcriptase (RT-) controls were used to check for DNA contamination. Quantitative PCR was performed in technical triplicates using TaqMan™ Gene Expression Master Mix (#4369016, Thermo Fisher Scientific) on a QuantStudio 7 Pro Real-Time PCR system (Applied Biosystems). Two housekeeping genes were used for normalization: RPL13A (Hs04194366_g1) and CETN2 (Hs00942570_g1). Additional probes used included: LGR5 (Hs00969422_m1), KRT20 (Hs00300643_m1), and GABRA2 (Hs00168069_m1). All probes were purchased from Thermo Fisher Scientific. For analysis, the comparative Ct method was used, with ΔCt values calculated relative to each housekeeping gene. ΔΔCt values were then calculated relative to the 7 dps sample. Expression levels are presented as log2 of 2^-ΔΔCt.

### The Cancer Genome Atlas (TCGA) and Human Protein Atlas tumors expression analysis

Expression data from different tumor types were obtained courtesy of the Human Protein Atlas (www.proteinatlas.org), consisting on RNA expression based on 8384 samples from 31 prognostic (TCGA) and validation cancers corresponding to 21 cancer types (38–40). The data is based on The Human Protein Atlas version 25.0 and Ensembl version 109. GABRA2 RNA expression is shown as protein-coding transcripts per million (pTPM). Box plots represent the median (line) and the interquartile range (25^th^-75^th^ percentiles), with whiskers extending to 1.5 times the interquartile range. Outliers are not shown.

### Flow cytometry analyses

Organoids were collected by centrifugation and dissociated into single cells using TrypLE Express (#12605-010, Gibco). For cytokine experiments, organoids were cultured for 11 days, after which human IL-6 (50 ng/mL; #130-093-931, Miltenyi Biotec) or human recombinant TGF-β1 (5 ng/mL; #100-21-10UG, PeproTech®) was added for 3 days prior to collection and digestion. Cells were then incubated with a primary antibody (anti-GABRA2, 1:100; #BS-12061R, Thermo Fisher Scientific) for 1 h in 0.5% BSA (#130-091-376, Miltenyi Biotec) PBS-EDTA (#130-091-222, Miltenyi Biotec), followed by a secondary antibody (anti-rabbit Alexa Fluor 488; #A21206, Invitrogen) for 30 min. DAPI was used as a viability dye, and cells were passed through a 50 μm CellTrics® filter (#04-0042-2317, Sysmex) prior to cytometric analysis to remove large clusters. Cytometry was performed on either a FACSymphony™ A1 (BD Biosciences) or a CytoFLEX LX (Beckman Coulter). Forward and side scatter plots were used to gate single cells, and DAPI positivity was used to exclude dead cells from analysis. Samples without primary antibody staining were used to set the positive gate for target protein expression. FlowJo™ v10 Software was used for analysis.

### Enzyme-Linked Immunosorbent Assay (ELISA)

Conditioned media (CM) from organoid cultures was collected at the indicated time points, centrifuged at 450 x g for 10 min at 4°C to remove cell debris, and the supernatant was collected and stored at −80°C until use. ELISA experiments of CM were performed in technical duplicates against IL-6 (ELH-IL6, RayBiotech), IL-6 sR (ELH-IL6sR, RayBiotech), and TFG-β1 (ELH-TGFb1, RayBiotech), following the kit’s recommendations. For TGF-β1 detection, samples were activated before testing according to the manufacturer’s instructions. Absorbance was measured at 450 nm using a GloMax® Discover microplate reader (Promega). Cytokine concentrations were determined using standard curves, and background signal was subtracted using a blank control (complete growth medium).

### Bulk RNA sequencing (bulk RNA-seq)

Organoids were cultured for 11 days, after which human IL-6 (50 ng/mL; #130-093-931, Miltenyi Biotec) or human recombinant TGF-β1 (5 ng/mL; #100-21-10UG, PeproTech®) was added for 3 days prior to collection. RNA was extracted using the Maxwell® RSC miRNA Tissue Kit (#AS1460, Promega), according to the manufacturer’s protocol. RNA was quantified using the Qubit™ RNA BR Assay Kits (Thermo Fisher Scientific) and quality evaluated using the Agilent RNA 6000 Nano Kit (Agilent Technologies, Santa Clara, CA, USA). Three independent experiments were performed and analyzed together. Gene expression libraries were obtained starting from 500 ng of RNA using TruSeq Stranded mRNA Library Preparation Kit (Illumina Inc., USA), following the manufacturer’s instructions. Final libraries were quantified using the Qubit dsDNA HS Assay Kit (Thermo Fisher Scientific), and their fragment distribution was evaluated using the DNA 1000 Kit Assay Kit (Agilent Technologies, Santa Clara, CA, USA). Equal amounts of DNA libraries were pooled and sequenced on the NovaSeq 6000 sequencer (Illumina Inc., USA). Sequencing settings were: read 1: 150 bp; read 2: 150 bp; dual index: 8 bp. *Quality control, alignment, and expression quantification*: Sequencing quality was initially assessed using FastQC (v0.11.9), and results were summarized with MultiQC (v1.14). Reads were aligned using STAR (41) (v2.7.1a; parameters: – outSAMunmapped Within –outFilterMultimapNmax 10 –outFilterMultimapScoreRange 3 – outFilterMismatchNmax 999 –outFilterMismatchNoverLmax 0.04) against the human reference genome (GRCh38.p10, GENCODE v27). All samples yielded more than 30M reads, with ≥ 65% of reads assigned to annotated genes, and therefore met the quality criteria for inclusion in downstream analyses. The computational pipeline used for these analyses is available at: https://github.com/molinerisLab/StromaDistiller. *Differential expression analysis*: Differential gene expression analysis between untreated and IL-6 or TGF-β1-treated samples was performed using DESeq2 (v1.38.3) within the R environment (v4.2.2). The model was specified using the design formula ∼ condition + replicate, where condition represents the culture condition (with IL-6 or TGF-β1 or without any addition) and replicate accounts for the three biological replicates. Prior to testing, lowly expressed genes were filtered out by removing those with fewer than 5 reads in more than one sample. Differentially expressed genes (DEGs) were identified using an absolute LFC ≥ 0.5849625 (corresponding to a ≥ 1.5-fold change) and an adjusted of *p*-value < 0.05. The resulting DEG lists (Supplementary Table 3-4) were used for Gene Set Enrichment Analysis (GSEA) using the Python package gseapy (v1.1.4). Enrichment analysis was performed in preranked mode on the gene ranking file derived from differential expression statistics. Gene sets from the Molecular Signatures Database (MSigDB, v2023.2) were used, including Hallmark and canonical pathway collections (C2, C5, C6, C8).

Graphical representations of the GSEA results were generated by plotting normalized enrichment scores (NES) for significantly enriched pathways (adjusted *p*-value < 0.05). Final plots were obtained by highlighting a selected subset of biologically relevant terms.

### Organoid-initiation assay

Organoids were cultured for 11 days in 12-well plates, after which human IL-6 (50 ng/mL; #130-093-931, Miltenyi Biotec) or human recombinant TGF-β1 (5 ng/mL; #100-21-10UG, PeproTech®) was added in triplicate per condition for 3 days prior to collection and dissociation into single cells. Viable cells were counted using Trypan Blue (Invitrogen) to exclude dead cells, and the number of live cells was normalized to the mean of the control (CTL) samples to assess the effect of cytokine treatment on viability. Then, 1000 live cells per replicate were replated in a 48-well plate pre-coated with a 1:1 mixture of Matrigel® and growth medium, in technical triplicates, and cultured in 2% Matrigel® complete growth medium without cytokines. Cells were allowed to grow for 7 days, after which they were fixed with 4% PFA (sc-281692, ChemCruz), and stained with DAPI. Fluorescence and bright-field images of entire wells were acquired as z-stacks using a wide-field microscope Nikon Ti2 (Lipsi) with a Plan Apo 2X 0.1 NA objective as a stitched tile of two images per well. Projections of z-stacks were computed as maximum intensity or extended-depth of focus. Ilastik (version 1.3.3post3) software was used to segment individual organoids (42), and Fiji software was used to count the total number of organoids per well, applying a size threshold of 20-2500 μm (37). For comparisons, the total number of organoids was normalized to the mean of the CTL samples. Protocol scheme was created with BioRender.com.

### Viability assay

Organoid cells were plated on 96-well plates pre-coated with a 1:1 mixture of Matrigel® and growth medium and cultured for 7 days in 2% Matrigel® complete growth medium. Then, Oxaliplatin (CRC0322: 5μM; CRC1502: 3μM) and cytokines (human IL-6: 50 ng/mL; #130-093-931, Miltenyi Biotec; or human recombinant TGF-β1: 5 ng/mL; #100-21-10UG, PeproTech®) were added, in six replicates per condition, in 2% Matrigel® complete growth medium. Organoids were cultured for an additional 7 days, after which viability was assessed using the MTS assay (CellTiter 96® AQueous One Solution Cell Proliferation Assay, #G3581, Promega), following the kit’s protocol. Absorbance was measured at 490 nm using a GloMax® Discover microplate reader (Promega). Blank wells (growth medium only) were used to subtract background signal. Values were normalized to the untreated control (CTL) condition. Three independent experiments were performed per organoid line.

### Statistical analyses

Statistical analyses were performed using GraphPad Prism 10 (GraphPad Software, CA, USA), unless otherwise stated. Statistical significance was assessed using one-way ANOVA followed by Dunnett’s multiple comparisons test, unless otherwise stated. Normality and homoscedasticity were assumed for the data. When the assumption of homoscedasticity was not met, a Brown-Forsythe ANOVA followed by Dunnett’s T3 multiple comparisons test was used instead. For smFISH analyses, statistical significance of differences in the percentages of LGR5- and KRT20-positive cells were assessed using PERMANOVA. For RT-qPCR analyses, statistical significance was determined using a two-tailed one-sample *t*-test, with *p*-values corrected by Bonferroni adjustment. For flow cytometry analyses percentages were transformed o log_2_ of the fold change relative to the control, and the statistical significance was determined using a two-tailed one-sample t-test, with p-values corrected by Bonferroni adjustment. For bulk RNA-seq, statistical significance of the genes belonging to Stem or differentiated modules, overlapping with differentially expressed genes in bulk was assessed by a hypergeometric test, considering as background the 19,344 genes expressed in the bulk. A *p*-value of < 0.05 was considered statistically significant.

## RESULTS

### CRC organoids show transcriptionally definite stem-like and differentiated-like populations

In order to investigate the presence of transcriptionally and phenotypically distinct subpopulations in CRC, we performed scRNAseq across three CRC organoid lines with different mutational backgrounds. These included CRC0322 (APC^LOF^ / KRAS^WT^), CRC0327 (APC^WT^ / KRAS^WT^), and CRC1502 (APC^LOF^ / KRAS^G12D^) (Fig. 1, Supplementary Fig.1). We first considered organoids at 11 days-post-seeding (dps) (Fig. 1A). As expected, scRNAseq datasets revealed that cells were distributed across different phases of the cell cycle, confirming significant proliferative activity in all three models. To characterize populations, we started by identifying groups of genes with highly correlated expression (modules) by using Hotspot for each organoid line (Supplementary Fig. 1A) (34). We found that, beyond cell cycle phase-specific variations, the datasets were robustly captured by two principal, independent gene modules: one characterized by expression of stemness-associated genes (“Stem”) and another associated with differentiation (“Differentiated”) (Fig. 1A), reflecting a hierarchical cellular organization in the organoids, consistent with previous reports (4, 9). Notably, gene-modules were found to be overlapping across the different organoid lines (Supplementary Fig. 1B). In particular, the Stem modules were characterized by the expression of genes such as LGR5, CDCA7, ASCL2, MEX3A, and EPHB2, whereas the Differentiated modules displayed expression of markers including KRT20, TFF3, EMP1 and CEACAM7, among others (Supplementary Fig. 1C, Supplementary Table 1). We then applied module-specific clustering and identified cells most strongly associated with each transcriptional program based on module metagene expression. This enabled us to classify cells as Stem, Differentiated, double-positive, or double-negative, referred to as “Calls” (Fig. 1A). The identification of subpopulations associated to the two modules allowed us to isolate and characterize cells from each subpopulation in each dataset. Consistent with their respective Stem and Differentiated states, cells in the stem-like subpopulation were distributed across all phases of the cell cycle, whereas differentiated cells were predominantly in G1, consistent with reduced or absent proliferative activity (Fig. 1B).

**Figure 1:**
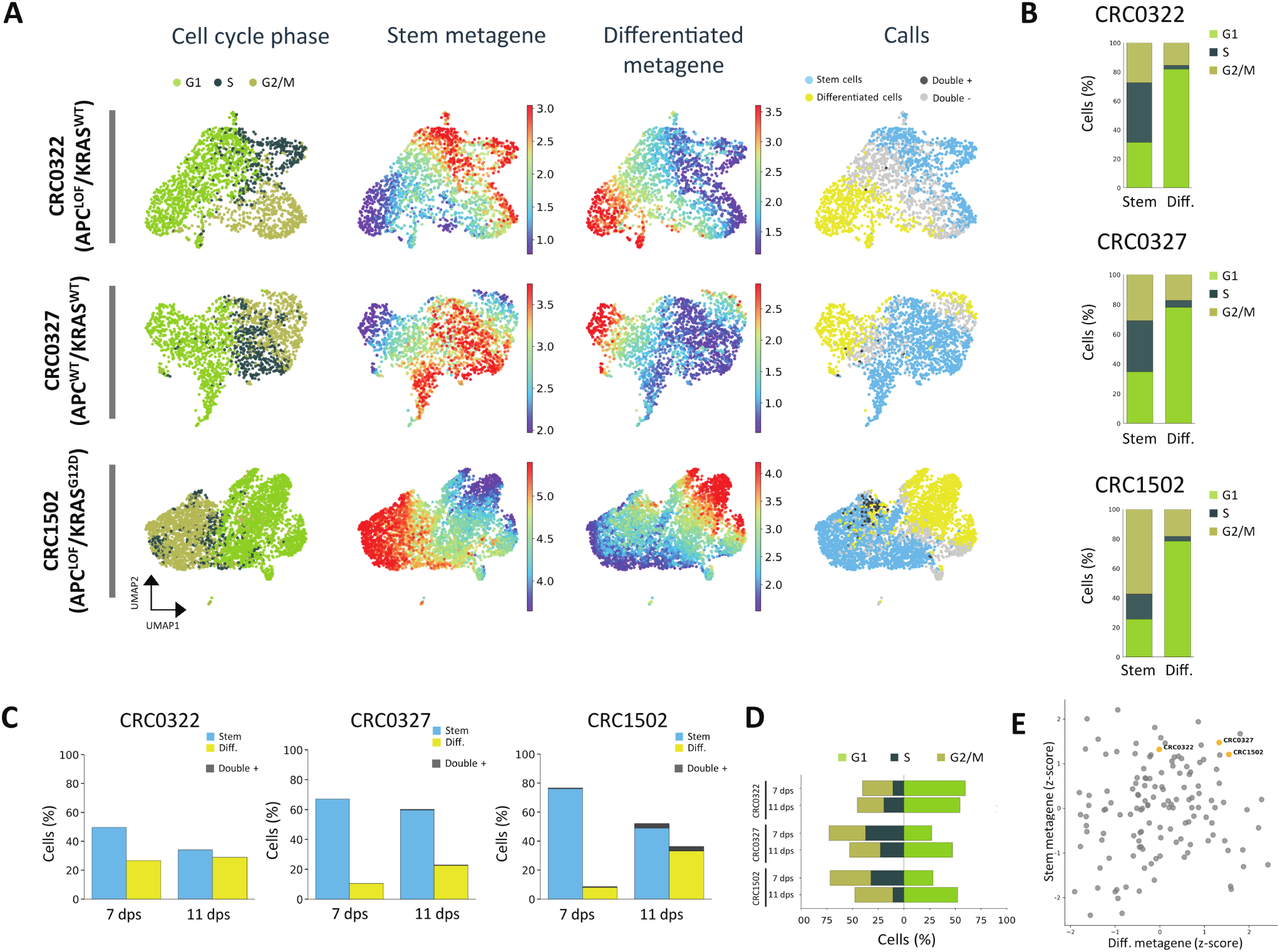
scRNAseq of CRC organoids. **A** Uniform Manifold Approximation and Projections (UMAPs) of CRC0322, CRC0327 and CRC1502 of cells 11 dps showing cell cycle phases, Stem module metagene expression, Differentiated module metagene expression, and the “calls” where cells were assigned to a subpopulation. **B** Quantification of the percentage of cells within each cell cycle phase, comparing stem-like cells (Stem) versus differentiated cells (Diff.) at 11 dps. **C** Histograms show the comparison of scRNAseq of organoids 7 dps versus organoids 11 dps. The calls were used to quantify the percentage of cells from the stem-like subpopulation (Stem) versus the cells from the differentiated-like subpopulation (Diff.) **D** Cell cycle phase comparison between organoids 7 dps and 11 dps. **E** Relative expression of the minimal modules metagenes from 11 dps samples for the Stem and the Differentiated (Diff.) subpopulations of CRC organoids from the XENTURION biobank. dps: days-post-seeding.

Next, to investigate the early dynamics of these subpopulations, we performed scRNAseq on the same organoid lines at 7 dps (Supplementary Fig. 1D). Using the same approach as before, we applied Hotspot to identify the gene modules and cell types present at this time point. As before, we identified both a stem-like subpopulation and a differentiated-like subpopulation (Supplementary Fig. 1D, Supplementary Table 1). The Stem and Differentiated gene modules for each organoid line showed partial overlap across time points (Supplementary Fig. 1E-G). We also observed temporal modulation of gene expression within these clusters. For example, the differentiated module at 7 dps included genes that were no longer expressed at 11 dps, and vice versa, indicating a dynamic acquisition of the phenotype (Supplementary Fig. 1E-G). Similar temporal dynamics were observed for the Stem modules (Supplementary Fig. 1E-G). We then compared the proportion of these two subpopulations within each time point and found that, overall, organoids appeared to shift toward a more differentiated state with time, accompanied by a reduction in the stem-like compartment (Fig. 1C). Cell cycle analysis revealed a shift toward G1 phase and a corresponding decrease in G2/M and S phase in the overall cell population of CRC0327 and CRC1502 at 11 dps, indicating that organoids become less proliferative as they grow (Fig. 1D), consistent with the decreasing stem-to-differentiated cells ratio. For CRC0322, no clear differences in the cell cycle phases were observed, which could reflect differences in proliferation or developmental rates between lines, with some organoids reaching a more stable cellular composition faster than others.

The three organoid lines tested exhibited a similar hierarchical structure, despite their different mutational backgrounds. This suggests that these subpopulations are not unique to our specific models. To assess whether this hierarchical organization is widespread across other samples, we examined the expression of minimal Stem and Differentiated modules, generated by intersecting the genes identified in each module across the three organoid lines at 11 dps, across the full set of organoids from the XENTURION biobank (Fig. 1E, Supplementary Table 2) (31). An overall percentage of 21% of organoids were found to express both minimal module metagenes (larger than the median for both and larger than 75th percentile for at least one), pointing at the presence of this hierarchical structure in multiple CRC models. All organoids considered in the manuscript were found in this set. In contrast, 22% of organoids were found to express neither module (smaller than the median for both and smaller than 25th percentile for at least one), suggesting that alternative population compositions may exist in CRC tumors.

### Single organoids are equipped with heterogeneously distributed LGR5+ and KRT20+ cell types reflecting diverse developmental trajectories

Although scRNAseq is a powerful technique for characterizing cellular transcriptional profiles, it requires disaggregation of organoids into single cells, resulting in the loss of spatial and structural information. In contrast, single-molecule Fluorescence In Situ Hybridization (smFISH) enables analysis of the transcriptional state of cells while preserving tissue architecture. In organoids, this approach allows us to determine whether different cell types coexist within the same organoid and to assess their organoid-level structure. We therefore applied this technique to identify stem-like and differentiated-like cells within CRC organoids, using LGR5 as a marker for stem-like cells and KRT20 for differentiated cells, both well-established markers for their respective subpopulations (4). In addition, we assessed the proliferative state of each cell using Ki67 (Fig. 2 and Supplementary Fig. 2). We analyzed the same three organoid lines, CRC0322 (Fig. 2A-D), CRC0327 (Fig. 2E-H) and CRC1502 (Fig. 2I-L), at 7 and 14 dps, extending the second time point beyond that used for scRNAseq to capture more pronounced differences. Anecdotally, we found that each organoid predominantly contained all three cell types, showing an intercalated pattern between them, rather than a clustered organization. We imaged 15-20 organoids per condition and quantified the number of spots of LGR5, KRT20 and Ki67 per cell (Fig. 2B, F, J, and Supplementary Fig. 2). This analysis revealed a clear divergence between cells expressing LGR5 and KRT20 at 14 dps, in contrast to the less distinct pattern observed at 7 dps (Fig. 2B, F, J).

**Figure 2:**
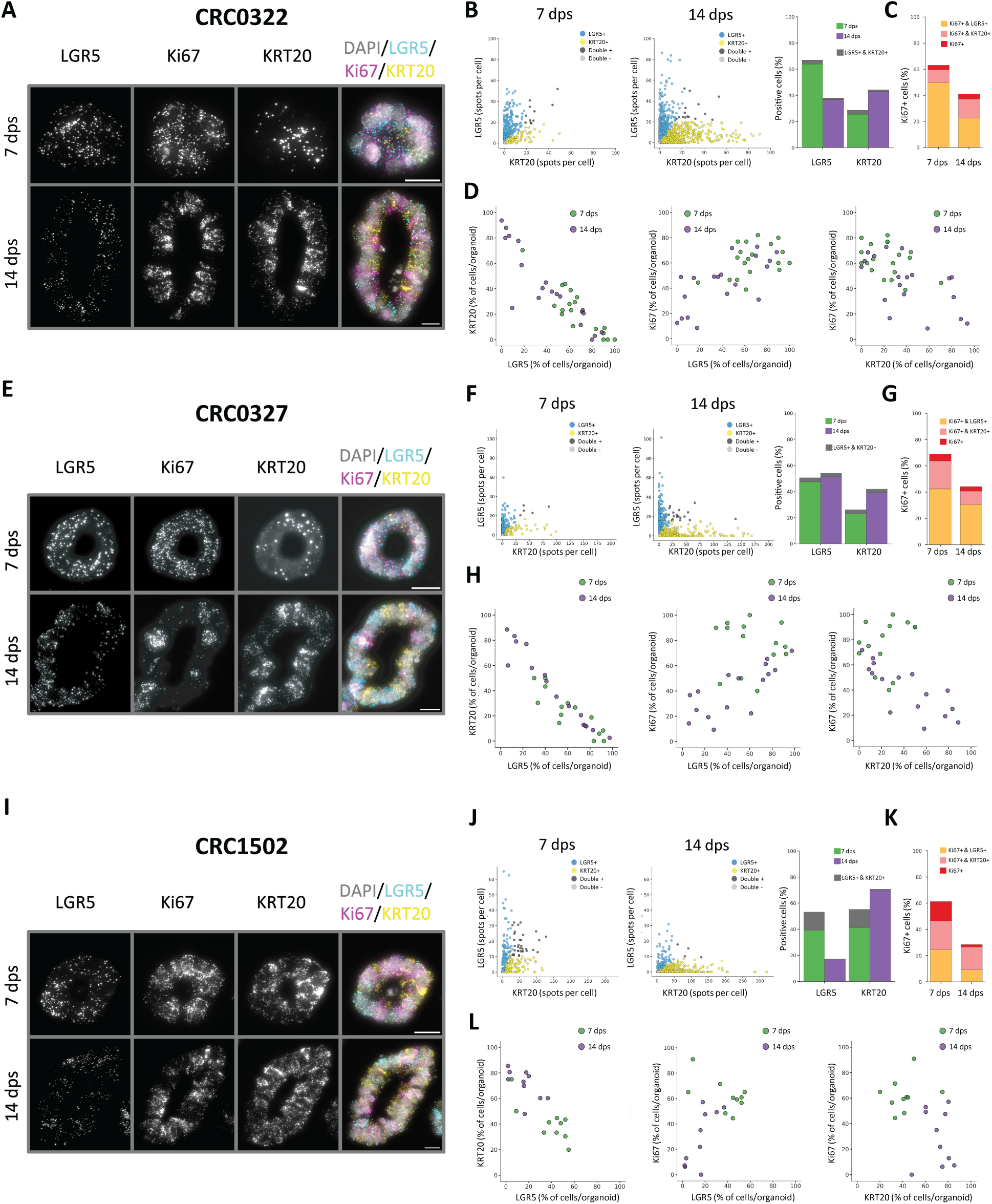
smFISH of LGR5, KRT20 and Ki67 in CRC organoids. **A-D** Representative fluorescent images of smFISH against LGR5, Ki67, KRT20, and DAPI of CRC0322 organoids 7 and 14 dps (A). Scatter plots show number of spots of LGR5 and KRT20 per cell (each dot represents a cell), histogram shows the percentage of positive LGR5 and KRT20 cells (B), and Ki67 positive cells (C) at 7 and 14 dps. Scatter plots show percentage of positive cells per organoid (each dot represents an organoid) (D). The median of (LGR5+;KRT20+) cells in organoids were (63.2;23.1)% (7d) vs (39.1;35.9)% (14d) (*p*-value = 0.0086, PERMANOVA). **E-H** Representative fluorescent images of smFISH targeted to LGR5, Ki67, KRT20, and DAPI of CRC0327 organoids 7 and 14 dps (E). Scatter plots show number of spots of LGR5 and KRT20 per cell (each dot represents a cell), histogram shows the percentage of positive LGR5 and KRT20 cells (F), and Ki67 positive cells (G) at 7 and 14 dps. Scatter plots show percentage of positive cells per organoid (each dot represents an organoid) (H). The median of (LGR5+;KRT20+) cells in organoids were (57.3;23.8)% (7d) vs (45.5;41.4)% (14d) (*p*-value = 0.207, PERMANOVA). **I-L** Representative fluorescent images of smFISH against LGR5, Ki67, KRT20, and DAPI of CRC1502 organoids 7 and 14 dps (I). Scatter plots show number of spots of LGR5 and KRT20 per cell (each dot represents a cell), histogram shows the percentage of positive LGR5 and KRT20 cells (J), and Ki67 positive cells (K) at 7 and 14 dps. Scatter plots show percentage of positive cells per organoid (each dot represents an organoid) (L). The median of (LGR5+;KRT20+) cells in organoids were (44.6;41.1)% (7d) vs (16.2;74.0)% (14d) (*p*-value = 0.0007, PERMANOVA). Fluorescent images: LGR5 (cyan), Ki67 (magenta), KRT20 (yellow), and DAPI (gray). dps: days-post-seeding. Scale bars = 20 μm.

Furthermore, the number of KRT20 spots per cells increased at 14dps with respect to 7dps across all organoid lines, supporting the notion that the differentiation status of these cells become more pronounced over time. We leveraged these data to classify each cell based on the expression of LGR5, KRT20, and Ki67 and quantified the percentage of cells at each time point. In all models, the proportion of KRT20-positive (KRT20+) cells increased at 14 dps, while CRC0322 and CRC1502 also showed a corresponding decrease in LGR5-positive (LGR5+) cells, as expected. Analysis of Ki67 showed fewer Ki67-positive (Ki67+) cells at 14 dps across all organoid lines, consistent with a decline in proliferative activity as the organoids developed (Fig. 2C, G, K). Lastly, we sought to assess differences between organoids by quantifying the proportion of each cell type within each organoid (Fig. 2D, H, L). Comparison across time points revealed that organoids at 7 dps were predominantly composed of LGR5+ and Ki67+ cells, while at 14 dps there was a reduction of these subpopulations and a concomitant increase in KRT20+ cells.

Surprisingly, we also observed marked inter-organoid heterogeneity, with some organoids exhibiting a high proportion of LGR5+ cells, while others showing a higher proportion of KRT20+ cells within the same time point and organoid line.

### A subset of differentiated cells is tracked by the expression of the GABRA2 protein

While transcriptional signatures allow characterization of different populations within a tumor, they lack the ability to fully resolve the longitudinal evolution of populations unless with transcriptional reporters, which are, however, experimentally challenging. Cell surface markers are instead powerful tools to follow the emergence and evolution of distinct cellular subpopulations. Thus far, we have used LGR5 and KRT20 as transcriptional markers for stem-like and differentiated cells, respectively. We next sought a cell surface marker capable of tracking changes in these subpopulations, with a particular focus on differentiation status. One candidate, emerging from our scRNAseq data and associated with all the gene modules for the differentiated subpopulations at both time points, was GABRA2, a subunit of the gamma-aminobutyric acid (GABA) channel typically expressed on the cell surface of neuronal and glial cells (Fig. 3A-C) (43). Interestingly, we found that, when measured using bulk gene expression assays during organoid development, GABRA2 expression was more indicative of population behavior than KRT20, showing a clear increase over time across all three models (Fig. 3D-F).

**Figure 3:**
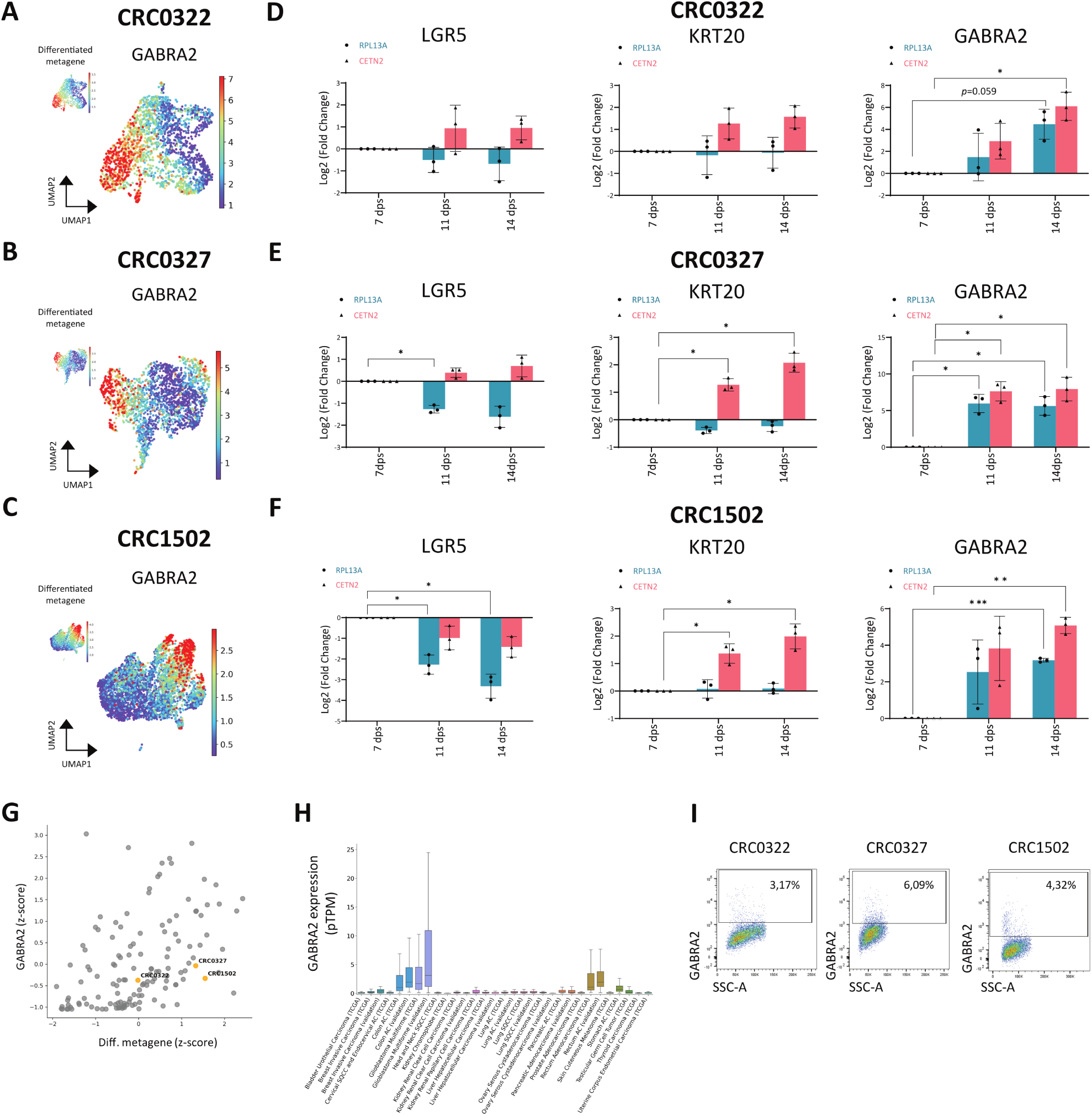
GABRA2 expression in CRC organoids. GABRA2 expression was analyzed in the scRNAseq 11 dps dataset (A-C) and by RT–qPCR (D-F). **A** CRC0322, **B** CRC0327, and **C** CRC1502 cells show GABRA2 expression within the Differentiated subpopulation. **D-F** Histograms show mean ± SD of RT-qPCR analysis of LGR5, KRT20, and GABRA2 levels over time of 3 independent experiments per organoid line. Expression is shown as log_2_ of the fold change relative to the 7 dps sample. Two housekeeping genes (RPL13 and CETN2) were used for normalization. Statistical significance was assessed using a one-sample t-test, with p-values corrected by Bonferroni adjustment. **G** Relative expression of GABRA2 versus the differentiated minimal module metagene (Diff.) minus GABRA2 of CRC organoids from the XENTURION biobank. **H** Expression of GABRA2 in samples from 21 tumor types from the TCGA and Human Protein Atlas (validation) dataset. Outliers are not shown. **I** Flow cytometry analysis confirms the presence of a fraction of cells with GABRA2 cell surface expression. dps: days-post-seeding; AC: Adenocarcinoma; SQCC: Squamous cell carcinoma; pTPM: protein-coding transcripts per million. \**p*<0.05; \*\**p*<0.01; \*\*\**p*<0.001.

Considering that GABRA2 is primarily considered a neuron-related protein, its detection in CRC cells was unexpected. We therefore examined the expression of GABRA2 transcript in the organoids from the XENTURION biobank, and related it to the expression of the previously defined minimal differentiated gene module (Fig. 3G and Supplementary Table 2). Not only was GABRA2 expressed across several organoid models, but its expression was also positively correlated with the differentiated metagene. We then analyzed GABRA2 expression in the TCGA and the Human Protein Atlas datasets across 21 tumor types to determine whether this finding was reproduced in other tumor types, and beyond the organoid model. Expression analysis revealed that, as expected, GABRA2 is detected in glioblastomas, and notably, it is also detected in colon and rectum adenocarcinomas (Fig. 3H).

We then used an antibody against GABRA2 to assess protein expression in our organoids and confirmed by flow cytometry of living cells that all three organoid lines contained a small fraction of GABRA2-positive (GABRA+) cells (Fig. 3I). The percentage of positive cells was lower than that detected by scRNAseq, a common observation when comparing transcript and protein levels, suggesting that we are not tracking the entire differentiated compartment, but rather likely capturing only a subset of these cells.

GABRA2 is one of the subunits that form the GABA heteropentameric channel (44). To determine whether other subunits were also expressed in our cells, and thus whether a functional GABA channel could be present, we examined the expression of all subunits in our scRNAseq dataset. We found that most subunits were either undetected or expressed at very low levels, with the exception of *GABRE* (Supplementary Fig. 3). However, the presence of only these two subunits would be insufficient to assemble a functional channel, leading us to conclude that the GABA channel is likely not present in these cells.

### The fraction of GABRA2+ cells can be modulated by exogenous cytokines

Cytokines represent key microenvironmental signals capable of reshaping cellular transcriptional states. Thus, we tested the effect of selected cytokines on organoid differentiation by flow cytometry using the GABRA2 antibody staining as a readout. The IL-6 receptor (IL6R) was among the genes associated within the differentiated subpopulation for CRC0322 and CRC0327, and was found expressed in some CRC1502 cells as well (Fig. 4A). Given that IL-6 is a cytokine commonly secreted by the tumor microenvironment, and that we also detected expression of its co-receptor gp130 (Supplementary Fig. 4), we evaluated its effects in our setup.

**Figure 4:**
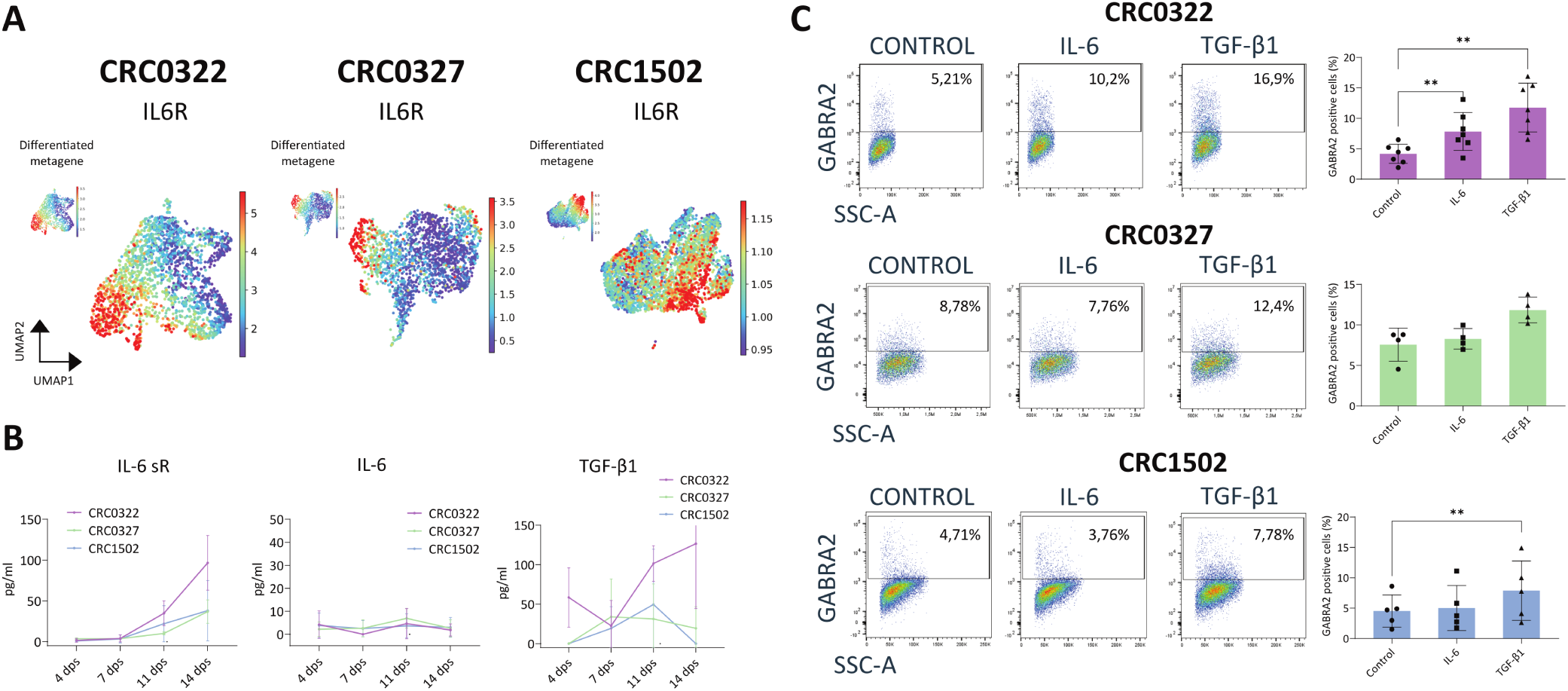
Cytokine modulation of differentiation. **A** scRNAseq analysis at 11 dps showing that CRC0322, CRC0327, and CRC1502 cells exhibit IL6R expression. **B** ELISA measuring the concentration of IL-6 sR, IL-6 and TGF-β1 in the conditioned media of CRC organoids over time. Curves show the average of 2 independent experiments. **C-E** Flow cytometry analysis of CRC0322 (C), CRC0327 (D), and CRC1502 (E) of organoids after 3 days of incubation with either IL-6 (50 ng/ml) or TGF-β1 (5 ng/ml). Histograms show the mean ± SD of the percentage of GABRA2 positive cells for 4-7 independent experiments. Statistical significances were analyzed by one-sample t-tests of the log_2_ fold change relative to the Control, with p-values corrected by Bonferroni adjustment. dps: days-post-seeding. \*\*\**p*<0.001.

In addition, we considered TGF-β1, another cytokine frequently present in the microenvironment, known to exert context-dependent effects on tumor cells (45). Our scRNAseq data confirmed the expression of TGF-β receptors in our cells, particularly of TGFBR1 (Supplementary Fig. 4). Furthermore, targeted sequencing of all three models revealed either wild-type profiles for TGF-β receptors and SMAD proteins, or very low variant allele frequencies (31).

We first analyzed the endogenous secretion of these cytokines by measuring their concentration in the conditioned media of organoids over time using ELISA, as well as the presence of the soluble IL-6 receptor (IL-6 sR), which has been reported to exist as a cleaved form, particularly in tumor contexts (46) (Fig. 4B). We observed increasing concentrations of IL-6 sR over time in all organoid lines, consistent with both the overall increase in cell number and with the increasing proportion of differentiated cells during organoid development. In contrast, IL-6 levels were nearly undetectable, whereas TGF-β1 showed variable concentrations over time, being either undetected or present at low levels.

Next, we treated organoids with either IL-6 (50 ng/mL) or TGF-β1 (5 ng/mL) for 3 days and performed FACS analysis using the GABRA2 antibody to assess shifts in differentiation. In CRC0322, TGF-β1 treatment induced a significant increase in the proportion of GABRA2+ cells, while IL-6 showed a similar, albeit weaker, trend (Fig. 4C). Conversely, CRC0327 and CRC1502 responded similarly to TGF-β1, though to a lesser extent, and showed no detectable response to IL-6 (Fig. 4D-E). The absence of an IL-6 response in these two lines, compared to CRC0322, is consistent with the lower levels of IL-6 sR detected in their conditioned media (Fig. 4B).

### TGF-β1 induces differentiation and enhances response to oxaliplatin in selected models

CRC0322 appeared to be the most responsive to both IL-6 and TGF-β1. We therefore used this model to further investigate the effect of these cytokines on differentiation. We first performed smFISH for LGR5, Ki67, and KRT20 on organoids treated for 3 days with either IL-6 or TGF-β1 (Fig. 5A-C, Supplementary Fig. 5A). IL-6 did not produce a clear effect on the total proportion of the different cell types. In contrast, TGF-β1 treatment led to a clear increase in KRT20+ cells, accompanied by a decrease in LGR5+ and Ki67+ cells (Fig. 5A-B). Consistently, analysis of cell-type composition at the organoid level showed that control (CTL) and IL-6-treated organoids exhibited similar profiles, whereas TGF-β1-treated organoids displayed a shift towards a higher proportion of KRT20+ cells (Fig. 5C). Furthermore, TGF-β1 appeared to induce a more pronounced expression of KRT20 at the single-cell level than that observed under basal conditions, as evidenced by a marked increase in the number of KRT20 spots per cell (Supplementary Fig. 5A).

**Figure 5:**
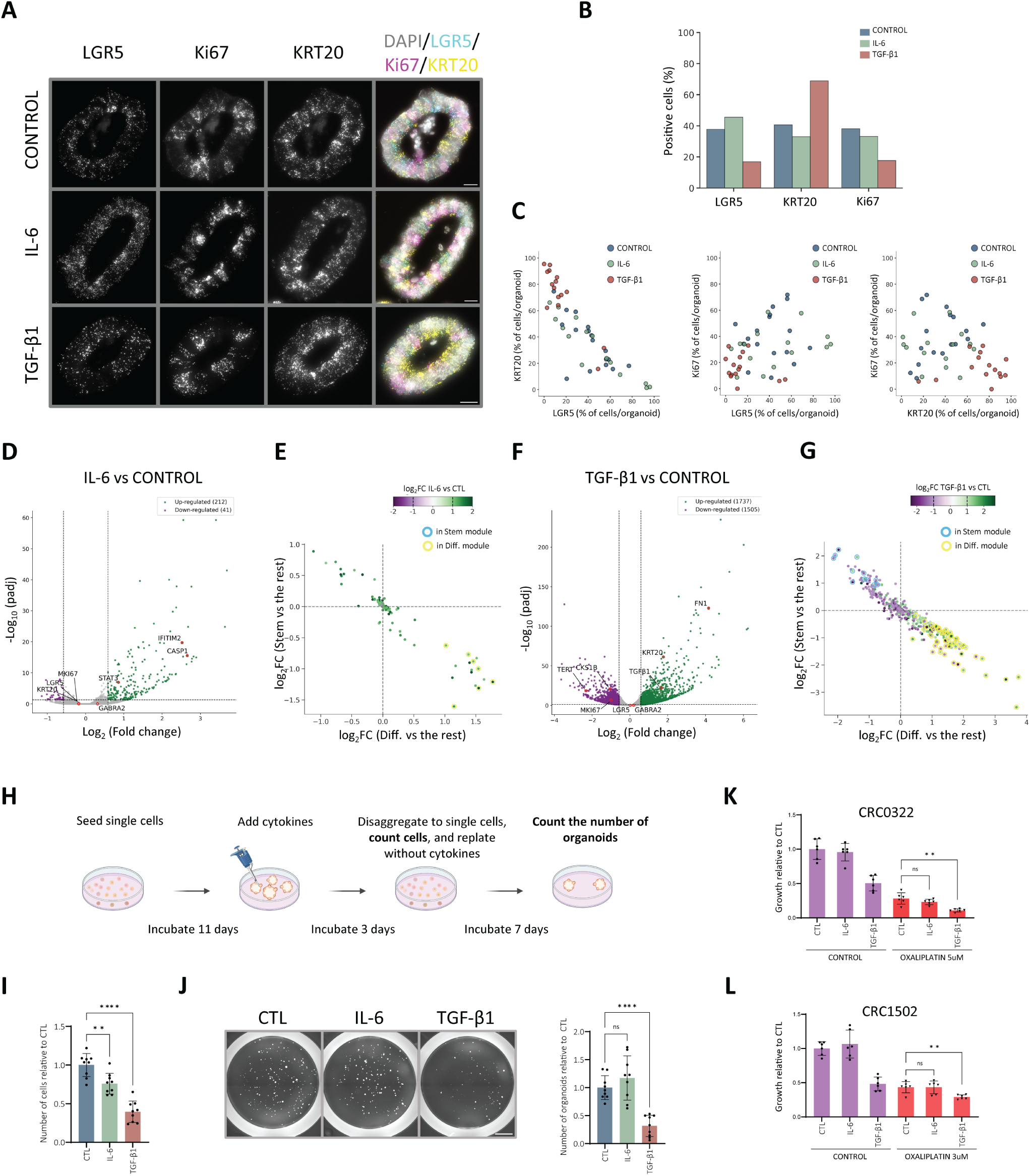
Effect of IL-6 and TGF-β1 on CRC organoids and response to chemotherapy. **A** smFISH of LGR5, KRT20 and Ki67 in CRC0322 after 3 days with IL-6 (50 ng/ml) or TGF-β1 (5 ng/ml). Scale bars = 20 μm. **B** Histogram shows the percentage of positive LGR5, KRT20 and Ki67 cells. **C** Scatter plots show percentage of positive cells per organoid (each dot represents an organoid). The median of (LGR5+;KRT20+) cells in organoids were (41.7;37.5)% CONTROL, (38.7;40.5)% IL-6, (10.7;72.3)% TGF-β1 (CONTROL vs IL-6 *p*-value = 0.7279, CONTROL vs TGF-β1 *p*-value = 0.0001, PERMANOVA test). **D, F** Volcano plots of bulk RNA-seq of CRC0322 after 3 days with IL-6 (D) or TGF-β1 (F). **E, G** Scatter plots show upregulated and downregulated genes (p<0.05, |log_2_FC|>1; color scale) for IL-6 (E) or TGF-β1 (G) treated organoids and log_2_FC in the scRNAseq 11 dps sample within cells classified as Stem or Differentiated versus all other cells. Genes belonging to either module are circled. **H** Scheme of the organoid-initiation assay. **I** Histogram shows the mean + SD of number of live cells relative to control (CTL) after IL-6 or TGF-β1 treatment of CRC0322 from 3 independent experiments pooled together (ANOVA followed by Dunett’s test versus CTL). **J** Images of organoids after replating 1000 live cells from previously treated organoids (scale bar = 2000 μm). Histogram shows the mean + SD of the number of organoids relative to the CTL from 3 independent experiments pooled together (ANOVA followed by Dunett’s test versus CTL). **K, L** Representative MTS viability assay of CRC0322 (K) and CRC 1502 (L) after 7 days with oxaliplatin plus IL-6 or TGF-β1 (replicates=6) (Brown-Forsythe ANOVA followed by Dunett’s T3 test (CRC0322); ANOVA followed by Dunett’s test (CRC1502) of oxaliplatin treated conditions versus oxalipatin CTL). Values were normalized to the untreated CTL. Experiments were repeated 3 times, obtaining similar results. Fluorescent images: LGR5 (cyan), Ki67 (magenta), KRT20 (yellow), and DAPI (gray). dps: days-post-seeding; ns: not significant; FC: Fold change. **p<0.01; ****p<0.0001.

Next, we assessed whether these changes were also evident at the bulk population level. To this end, we performed bulk RNA sequencing (bulk RNA-seq) on treated CRC0322 organoids (Fig. 5D-G, Supplementary Fig. 5B-C, Supplementary Table 3-4). Gene set enrichment analysis (GSEA) revealed that IL-6 treatment triggered a strong interferon response, primarily associated with the expression of IFITM2 and CASP1, as well as the expected activation of the IL-6/JAK/STAT3 pathway (Supplementary Fig. 5B, Supplementary Table 3). Treatment with IL-6 did not induce a clear increase of KRT20 or GABRA2, nor did it prompt a decrease in LGR5 and Ki67 (Fig. 5D). When comparing the differentially expressed genes in the bulk to the genes from the Stem and Differentiated modules of CRC0322 at 11 dps, we only observed 7 genes belonging to the Differentiated module that were upregulated (*p*-value = 2.77 × 10⁻⁴, hypergeometric test), while we did not find any Stem module gene downregulated, supporting the notion that no clear modulation of either module was induced (Fig. 5E). As for the treatment with TGF-β1, GSEA analysis revealed a strong enrichment of the TGF-β signalling pathway, including increased expression of TGFB1 itself, as well as activation of epithelial-to-mesenchymal transition (EMT) programs associated with genes such as Fibronectin 1 (FN1) (Supplementary Fig. 5C, Supplementary Table 4). Additionally, pathways involved in cell cycle progression were downregulated, consistent with decreased expression of genes such as CKS1B. Among other genes, TGF-β1 treatment induced a marked upregulation of KRT20, accompanied by a decrease in Ki67 expression (Fig. 5F).

Notably, LGR5 and GABRA2 did not show significant modulation, suggesting that some aspects of cellular heterogeneity may not be fully captured at the bulk level. Additionally, we observed downregulation of the telomerase subunit TERT, whose high expression is commonly associated with stem-like cells (47). Comparison of the differentially expressed genes with genes from the Stem and Differentiated modules revealed that 41 genes out of 212 from the Differentiated module were upregulated upon TGF-β1 treatment (*p*-value = 4.19 × 10⁻²¹, hypergeometric test), along with downregulation of 8 genes out of 68 from the Stem module, further supporting an effect of this cytokine on differentiation (*p*-value = 3.51 × 10⁻⁵, hypergeometric test) (Fig. 5G).

Our data support the notion that TGF-β1 increases the proportion of differentiated cells within organoids. Given that differentiated cells are generally less proliferative, we next investigated the functional consequences of cytokine treatment. To this end, we performed an organoid-initiation assay (Fig. 5H-J). First, CRC0322 organoids were treated with IL-6 or TGF-β1 for 3 days, after which they were dissociated into single cells and counted. This analysis revealed that TGF-β1 treatment significantly reduced the number of viable cells (Fig. 5I). This decrease may reflect apoptosis induction, as previously reported for TGF-β1 (48, 49), or also enhanced differentiation leading to a reduction in the fraction of proliferative cells. We then assessed the organoid-forming capacity by replating 1,000 pre-treated cells and counting the number of organoids formed after 7 days. Consistent with an increase in differentiated cells, TGF-β1 pre-treatment led to a significant reduction in organoid-forming efficiency (Fig. 5J), whereas IL-6 had no significant effect. Taken together, these results demonstrate that TGF-β1 induces functional changes in cell behavior, reflecting a change in population composition within organoids and a consequent change in cellular states rather than solely transcriptional alterations.

Lastly, we investigated the effect of cytokine stimulation on the response to oxaliplatin, a commonly used chemotherapeutic agent, in CRC0322 as well as CRC1502, which also showed a small induction of GABRA2+ cells in FACS analysis. We first performed dose-response assays for each organoid line to determine the appropriate concentration of oxaliplatin (Supplementary Fig. 5D-E). We then assessed organoid viability after 7 days of co-treatment with oxaliplatin in combination with either IL-6 or TGF-β1 in CRC0322 (Fig. 5K) and CRC1502 (Fig. 5L). IL-6 did not significantly affect the response to oxaliplatin, consistent with its limited impact on differentiation. In contrast, TGF-β1 in combination with oxaliplatin resulted in a significant reduction in viability compared to oxaliplatin treatment alone for both organoid lines. Overall, our results therefore suggest that promoting differentiation can enhance sensitivity to oxaliplatin.

## DISCUSSION

Single cell resolution study of intratumor phenotypic heterogeneity has advanced our understanding of key processes such as tumor development, progression, metastasis, and response to treatment (12, 50–52). Our results further support the concept that tumors comprise transcriptionally distinct subpopulations with different hierarchical states, behaviors, and functional potentials. Indeed, a subset of CRC organoids are composed of two subpopulations previously described: stem-like cells, associated with expression of genes such as LGR5, and differentiated cells, associated with KRT20 (4, 51), with a distinct lineage hierarchy, reminiscent of a normal multi-type tissue. Additional subpopulations involved in treatment-induced transitional states, therapy tolerance, and phenotypic plasticity have also been described, including cells with a fetal-like transcriptional signature associated with YAP activation (53–56), cells with a Paneth-like signature (14, 57), slow-cycling stem-like cells (15), and high-relapse cells, with slow-cycling stem-like cells and high-relapse cells associated with expression of the genes MEX3A and EMP1, respectively (both of which were included in our Stem and Differentiated modules respectively)(21, 51, 58, 59). Deeper investigations on the origin of drug-tolerant or high relapse cells are needed in order to better exploit cell plasticity as a therapeutic leverage.

Organoid-level analysis of our CRC organoids enabled us to examine the percentage of different cell types within each organoid and, interestingly, we observed a significant inter-organoid variability within the same organoid culture. The presence of organoids with markedly different cellular compositions - such as those enriched in LGR5+ cells versus others with a higher prevalence of KRT20+ cells - reinforce once more the notion that differentiation paths are characterized by a strong stochasticity. This also points to the idea that, especially during ‘development’, tumor organoids might be characterized by very dynamic and unstable states, which might also hide priming and commitment of cells towards a particular state. Furthermore, a concomitant interpretation could be that transcriptional cell states are rather a continuous spectrum of states with different transition probabilities, that could ultimately result in the large observed inter-organoid variability. In this context, the strong effect of TGF-β1 on differentiation was evident not only by the increase in the proportion of KRT20+ cells, but also in the increased KRT20 expression per cell, accompanied by a concomitant reduction in organoid-initiating capacity. Along this line, while it is generally assumed that organoids arise from stem-like cells, single cell studies have shown that not all LGR5+ cells are transcriptionally equal (15, 58). Moreover, it is possible that other cell types, including KRT20+ cells, retain the capacity to form organoids, consistent with evidence that some KRT20+ cells can reacquire stem-like properties (4). All in all, these observations call for more in-depth analysis and surely highlight organoids as a valuable model to study lineage hierarchies and phenotypic plasticity.

The structured differentiation hierarchies present in patient-derived organoids were instrumental for untangling the role of TGF-β1, which has been reported to exert opposite effects depending on tumor stage. In pre-malignant lesions, this cytokine has been shown to promote cell cycle arrest and apoptosis, consistent with our observations, whereas in late-stage tumors it was shown to act as a tumor promoter, particularly through its pro-invasive, immunosuppressive and desmoplastic role (49, 60–64). TGF-β1 inhibitors are indeed being evaluated as a therapeutic option in metastatic CRC mostly in combination with immunotherapy (64). Along this line, in our setting, TGF-β1 also induces the upregulation of the gene EMP1. As mentioned above, EMP1 has been linked to a subpopulation referred to as High Relapse Cells (HRCs) - a fraction of which also express KRT20 - which are associated with metastatic initiation, in particular EMP1^+^/KRT20^-^ cells and associated to immune exclusion (51). These data might indicate that TGF-β1 exposure actively remodels the functional potential of the differentiated populations. In contrast, in our cell autonomous settings, TGF-β1 caused an increase in differentiation markers, a decreased cell cycle activity, a decreased clonogenic potential and an increased potency of oxaliplatin (65, 66). These results would envisage an antitumor activity for TGF-β1. Therefore, if on the one side TGF-β1 is associated to a negative impact on tumor progression, its inhibition could reduce the efficacy of chemotherapy in oxaliplatin-treated patients (67).

## CONCLUSIONS

Overall, our experimental evidence show that organoids are a valuable model system to address cell-state dynamics during tumor growth and phenotypic plasticity by exploiting response to exogenous stimuli and therapeutic strategies. Our results further support the notion that even if metastatic, some tumors retain the ability to fully respond to TGF-β1 via the classic tumor cell-autonomous mechanism. For tumor samples that do not carry TGF-β1 pathway mutations, exposure to this cytokine could represent a potentially useful therapeutic option to be exploited in combination with other pharmacological approaches.

## DECLARATIONS

### Ethics approval and consent to participate

Organoids used in this study were originally derived from tumor samples obtained from patients undergoing liver metastasectomy at the Candiolo Cancer Institute, Candiolo, Torino, Italy; Ospedale Mauriziano Umberto I, Torino, Italy; and Grande Ospedale Metropolitano Niguarda, Milano, Italy. Written informed consent was obtained from all patients prior to sample collection. Samples were procured, and the study was conducted within the concluded observational trial “Prospective study for the determination of the molecular profile of resistance to antineoplastic treatments” - PROFILING protocol No. 001-IRCC-00IIS-10 - approved by the Ethics Committee of the Candiolo Cancer Institute FPO IRCCS (authorization v. 11.0, dated July 13, 2022), in accordance with the Declaration of Helsinki guidelines.

### Consent for publication

Not applicable

### Availability of data and materials

The scRNAseq and bulk RNA-seq datasets used and/or analysed during the current study are available from the corresponding author upon reasonable request. The datasets will be deposited in the European Genome-phenome Archive (EGA).

### Authors’ contributions

SJF, AP, AB, and LT conceptualized the study. SJF, IC and MP performed the experiments. LPi and EG performed computational processing and bioinformatic analyses. SJF, LPi, and AP interpreted the data. IC, MP, EG, FG, SB, LP, AB and LT contributed to discussion of the results. SJF and AP drafted the manuscript. All authors revised the manuscript.

## Acknowledgements

We thank Annamaria Gullà, MD and Marcello Turi, PhD for providing access to the FACSymphony™ A1 (BD Biosciences). We acknowledge the precious work of our institute’s facilities: Dr. Roberta Porporato on smFISH; Dr. Alice Bartolini and Dr. Daniela Cantarella on single cell and bulk RNAseq; Dr. Paola Bernabei and Dr. Denis Baev for flow cytometry; Dr. Barbara Martinoglio for RT-qPCR.

The results shown in Figure 3H are in part based upon data generated by the TCGA Research Network: https://www.cancer.gov/tcga.

## Declaration of AI-tools in the writing process

ChatGPT-5.5 and Gemini 3 were used during the drafting of this manuscript to assist with grammar, spelling and readability, and for coding in the computational analyses. All content was revised by authors, who take full responsibility for the final content.

## Supplementary Material

Supplementary Figures: PDF file containing supplementary figures 1 to 5

Supplementary Tables 1: XLSX file with list of genes corresponding to the modules from 7 dps and 11 dps

Supplementary Table 2: XLSX file with list of genes corresponding to the minimal modules

Supplementary Table 3: XLSX file with bulk RNA-seq results from organoids treated with IL-6 versus the control

Supplementary Table 4: XLSX file with bulk RNA-seq results from organoids treated with TGF-β1 versus the control

## LIST OF ABBREVIATIONS

CRC: Colorectal cancer
dps: days-post-seeding
FC: Fold change
FDR: False Discovery Rate
GABA: gamma-aminobutyric acid
IL-6 sR: soluble IL-6 receptor
ISCs: Intestinal stem cells
PDXs: Patient-derived xenografts
scRNAseq: single-cell RNA sequencing
smFISH: single-molecule Fluorescence In Situ Hybridization
UMAPs: Uniform Manifold Approximation and Projections

