## Supplementary Figures 1 to 5 for "TGF-β1-induced differentiation enhances chemotherapy response in metastatic colorectal cancer organoids"

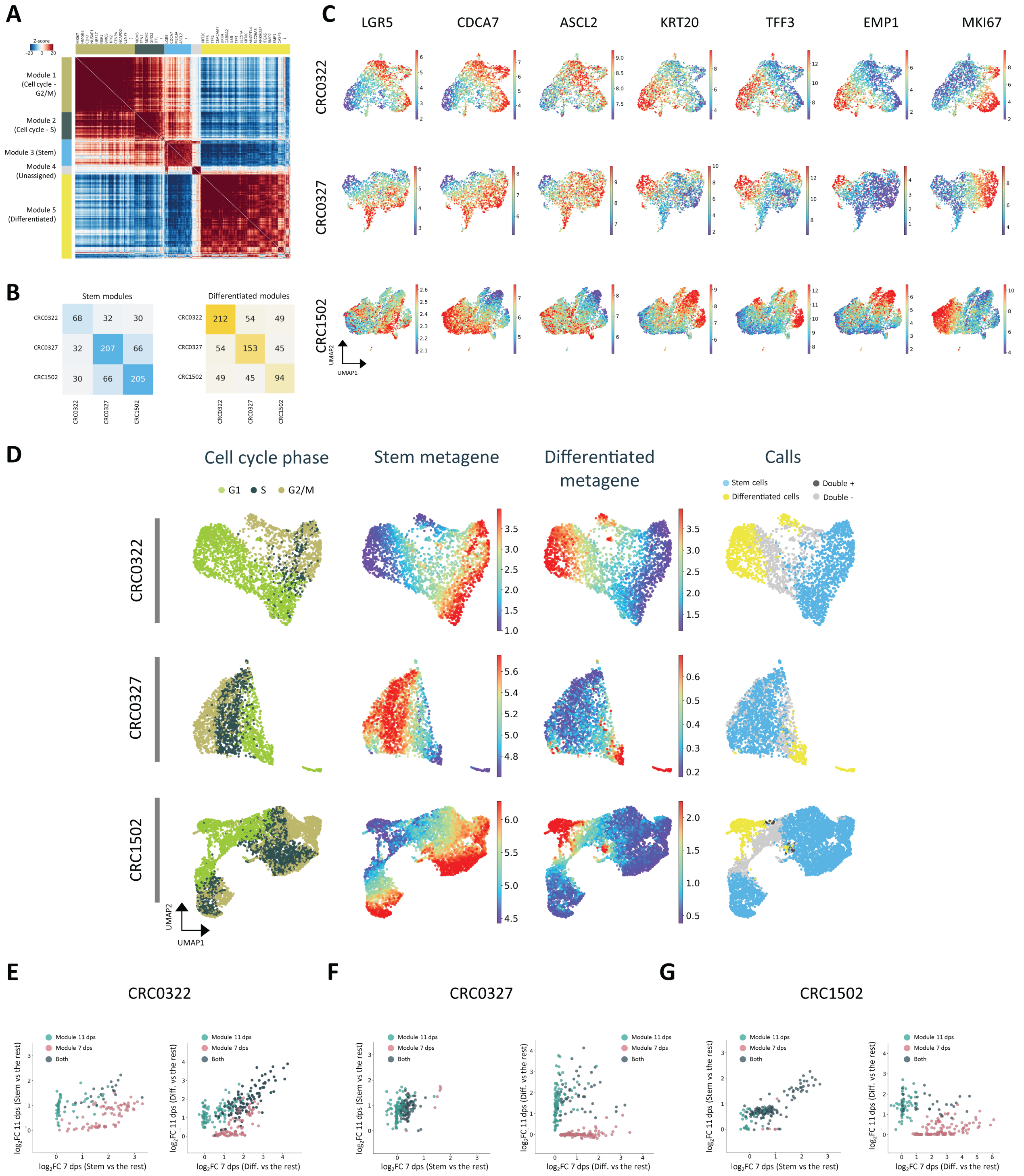


Supplementary Figure 1: scRNAseq of CRC organoids, clustering and 7 dps dataset. **A** Gene modules identified with Hotspot on CRC0322 at 11 dps used to define modules and perform subsequent clustering on cells. **B** Matrices showing common genes between Stem and Differentiated subpopulations across organoid lines at 11 dps. **C** UMAPs of selected genes associated to Stem and Differentiated modules as 11 dps. **D** UMAPs of scRNAseq of organoids at 7 dps showing cell cycle phase, expression of modules metagenes, and the "calls" where cells were assigned to a subpopulation. **E, F, G** Scatter plots of fold-change expression of all genes from the Stem (left) and Differentiated (right) modules corresponding to each time point, showing shared and unique genes between time points for CRC0322 (E), CRC0327 (F), and CRC1502 (G). dps: days-post-seeding; Diff.: differentiated; FC: Fold change.

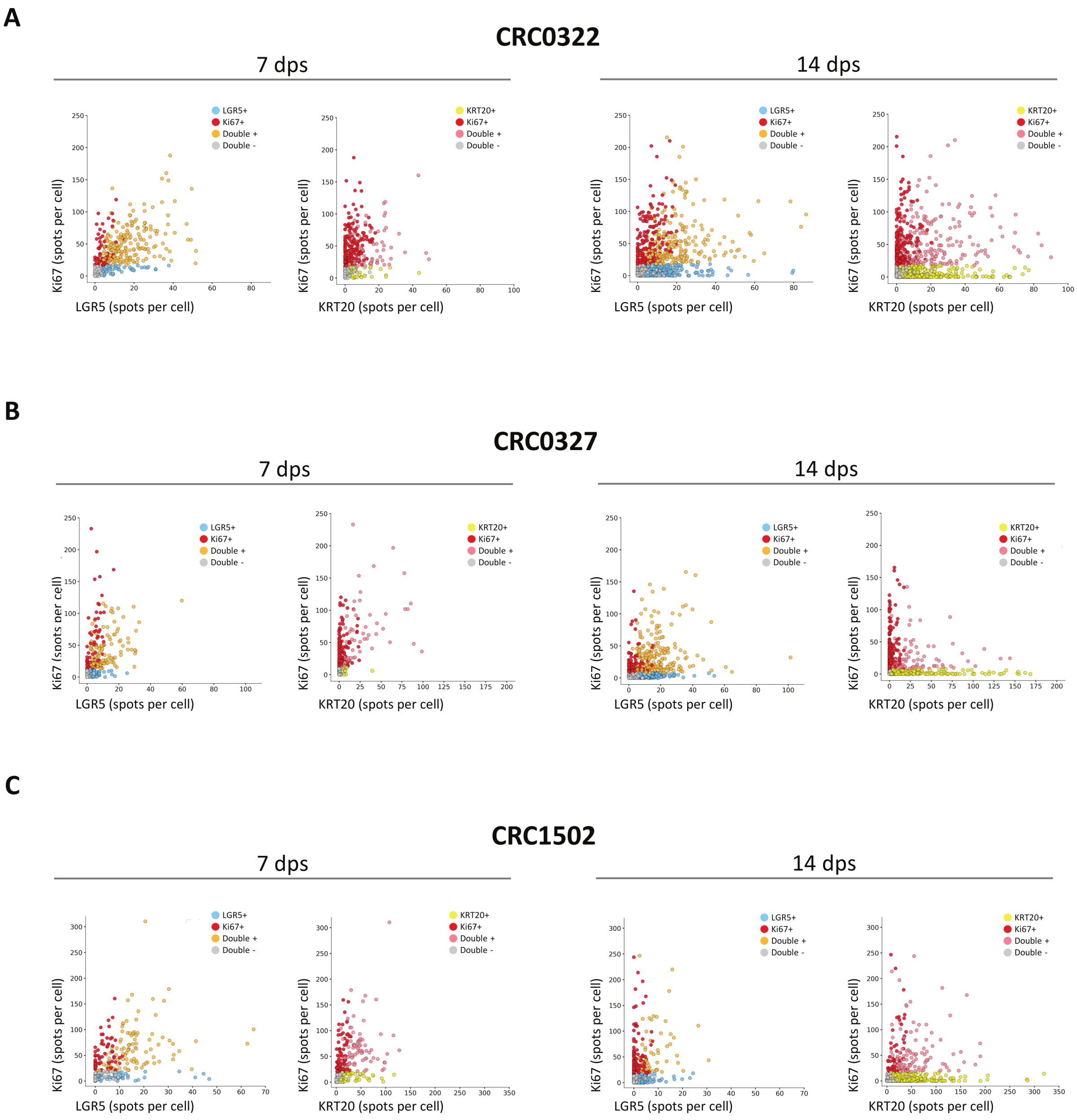


Supplementary Figure 2: smFISH of LGR5 and KRT20 versus Ki67 in CRC organoids at 7 and 14 dps. **A** Scatter plots show number of spots per cell of LGR5 versus Ki67 and KRT20 versus Ki67 of CRC0322 organoids. **B** Scatter plots show number of spots per cell of LGR5 versus Ki67 and KRT20 versus Ki67 of CRC0327 organoids. **C** Scatter plots show number of spots per cell of LGR5 versus Ki67 and KRT20 versus Ki67 of CRC1502 organoids. Each dot represents a cell. Quantifications of positive and negative cells for each gene are shown in Figure 2. dps: days-post-seeding.

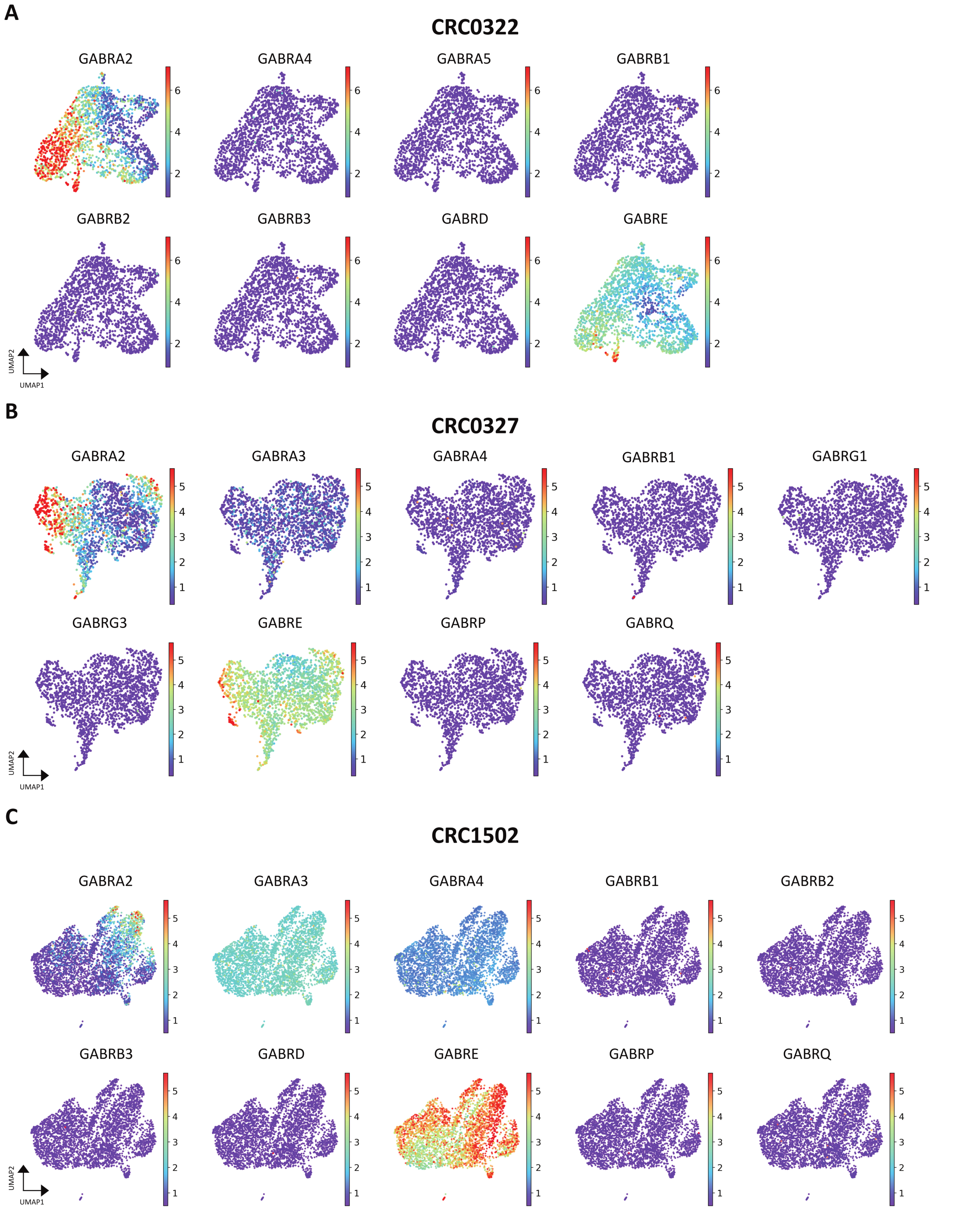


Supplementary Figure 3: Expression of subunits of the GABA channel in the scRNAseq 11 dps dataset. UMAPs from scRNAseq of the different subunits of the GABA channel found in CRC0322 (**A**), CRC0327 (**B**) and CRC1502 (**C**) organoids 11 dps. Scales were normalized for each organoids line for comparison purposes. Most subunits showed low or no expression and no colocalization with GABRA2, with the exception of GABRE. Undetected subunits are not shown. Lack of expression of other subunits suggests that the GABA channel might not be assembled. dps: days-post-seeding.

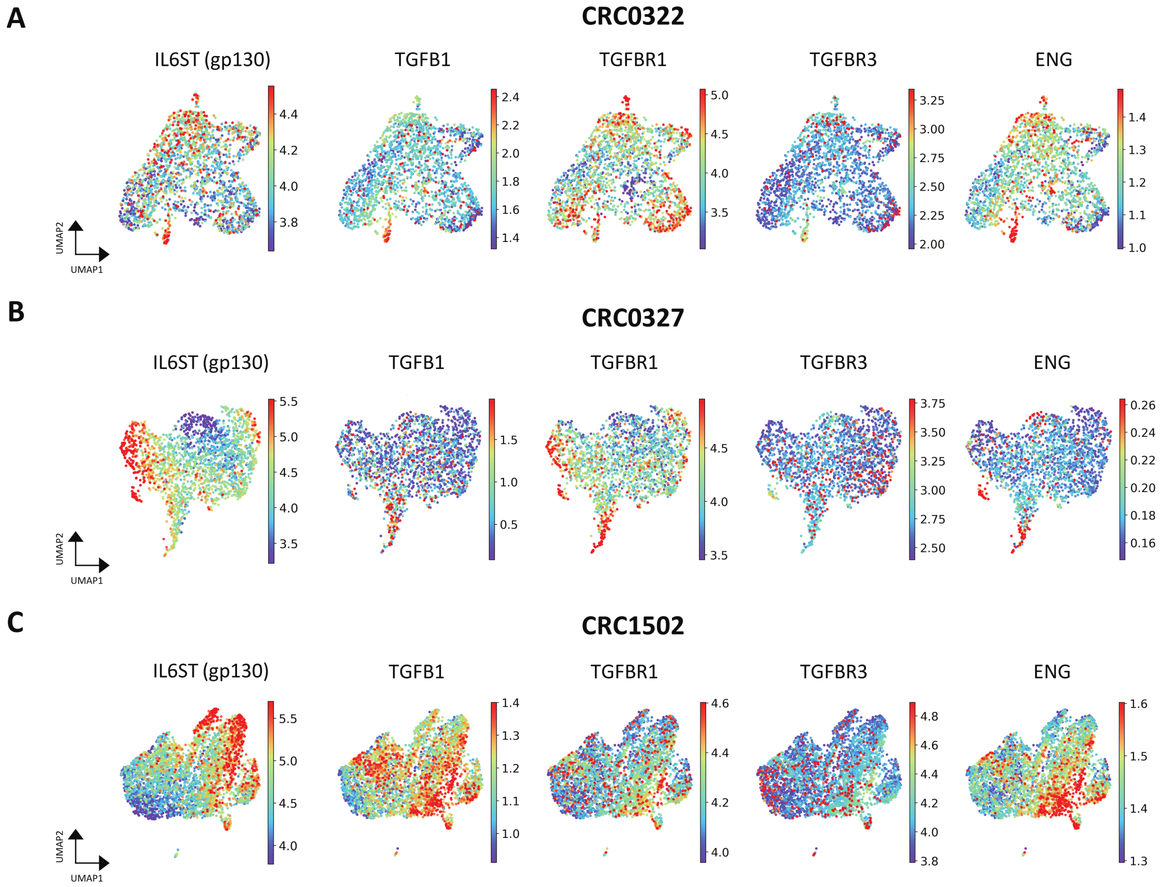


Supplementary Figure 4: Expression of co-receptor of IL-6, and TGF-β1 with its receptors in the scRNAseq 11 dps dataset. UMAPs from scRNAseq of the IL-6 co-receptor IL6ST (gp130), TGFB1, and TGF-β1 receptors in CRC0322 (**A**), CRC0327 (**B**) and CRC1502 (**C**) organoids at 11 dps. dps: days-post-seeding.

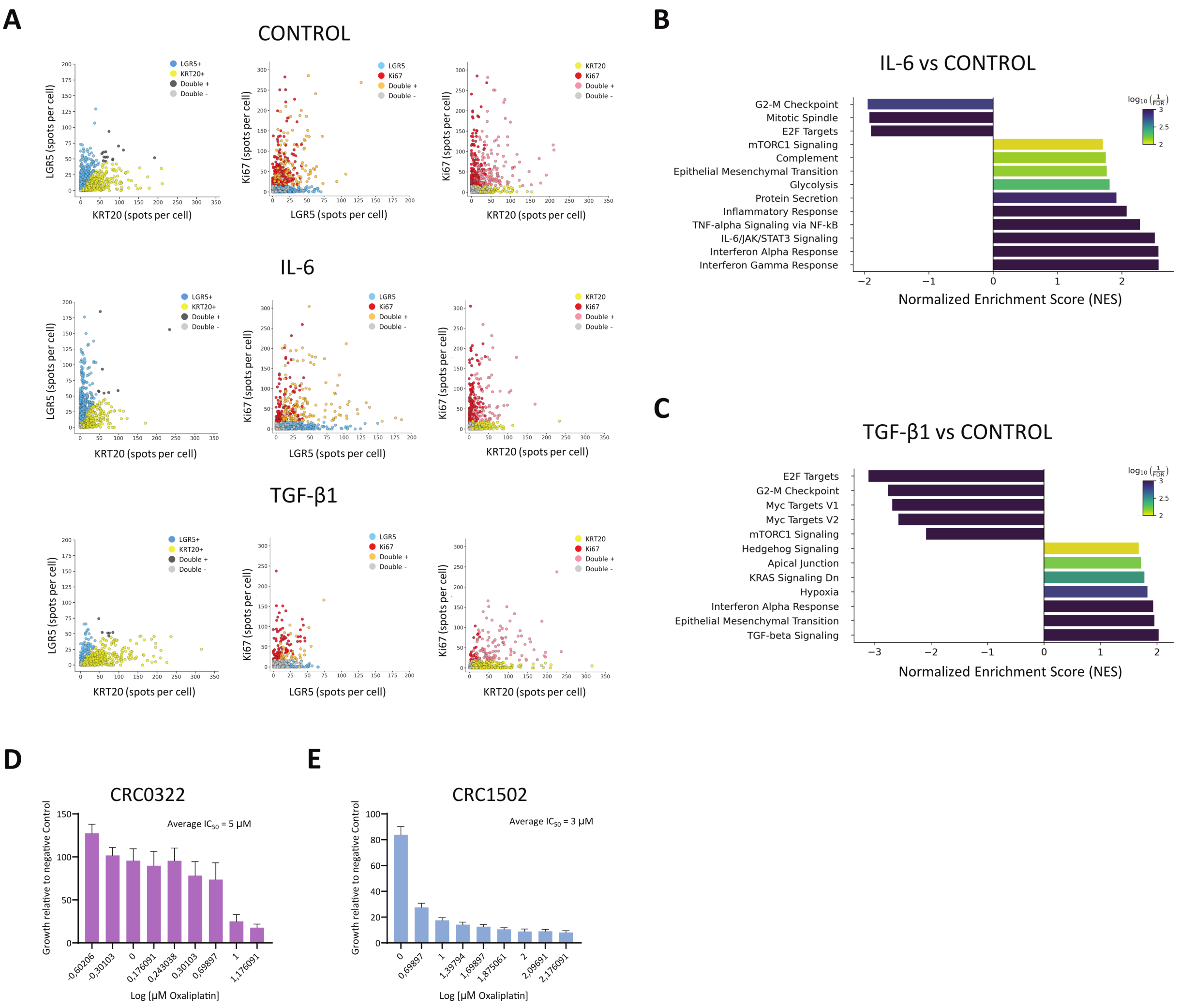


Supplementary Figure 5: smFISH of LGR5 and KRT20 and Ki67 in cytokine-treated organoids, and oxaliplatin dose-response curves. **A** Scatter plots show number of spots per cell of LGR5, Ki67 and KRT20 in CRC0322 organoids treated with either IL-6 (50 ng/ml) or TGF-β1 (5 ng/ml) or Control. Quantification of the number of cells called within each type per condition are depicted in Figure 5. **B, C** GSEA analysis of CRC0322 bulk RNA-seq data showing upregulated and downregulated pathways after IL-6 (B) or TGF-β1 (C) incubation. **D-E** Representative dose-response curves of CRC0322 (D) and CRC1502 (E) treated with oxaliplatin for 7 days. $IC_{50}$ values were calculated using nonlinear regression. Experiments were performed in duplicate for each line, and the selected $IC_{50}$ value represents the average of the two independent experiments. FDR: False Discovery Rate.
